# PlasmidHawk: Alignment-based Lab-of-Origin Prediction of Synthetic Plasmids

**DOI:** 10.1101/2020.05.22.110270

**Authors:** Qi Wang, Tian Rui Liu, R. A. Leo Elworth, Todd J Treangen

## Abstract

With advances in synthetic biology and genome engineering comes a heightened awareness of potential misuse related to biosafety concerns. A recent study employed machine learning to identify the lab-of-origin of DNA sequences to help mitigate some of these concerns. Despite their promising results, this deep learning based approach had limited accuracy, is computationally expensive to train, and wasn’t able to provide the precise features that were used in its predictions. To address these shortcomings, we have developed *PlasmidHawk* for lab-of-origin prediction. Compared to a machine learning approach, *PlasmidHawk* has higher prediction accuracy; *PlasmidHawk* can successfully predict unknown sequences’ depositing labs 63% of the time and 80% of the time the correct lab is in the top 10 candidates. In addition, *PlasmidHawk* can precisely single out the signature sub-sequences that are responsible for the lab-of-origin detection. In summary, *PlasmidHawk* represents a novel, explainable, accurate tool for lab-of-origin prediction of synthetic plasmid sequences. *PlasmidHawk* is available at https://gitlab.com/treangenlab/plasmidhawk.git

## Introduction

Thanks to the advancement of genome engineering and sequencing technology, researchers now have the capability to readily read and write DNA sequences^1^. This new technology has the promise of significantly improving the quality of human life through various fields, such as increasing agricultural yields^2^, accelerating drug discovery^3^ or advancing gene therapy^4^. While the use cases of this exciting technology enabling the bio-economy are largely positive, biosecurity, IP infringement, and potential misuse remain as concerns^5^. As a proof of principle, in response to previously outlined concerns, Allen *et al*. utilized a set of signature sequences with length k, also referred as signature k-mers, to differentiate artificial sequences from natural genomes and plasmids^6^. Although this promising k-mer based matching approach offers the ability to distinguish artificial sequences from a set of background sequences, there is still a need to develop a predictive pipeline that enables to handle an enormous amount of input sequences and reveal finer details of a given synthetic sequence. To meet the need, Nielsen *et al*. introduced a software tool to link artificial sequences with their depositing labs by using deep learning^7^. Despite the complex computational challenge, the prediction accuracy was promising: 48% accuracy in correctly identifying the lab-of-origin for an unknown sequence if allowed one prediction, and up to 70% accuracy if allowed ten predictions. To date, deep learning has been wildly applied in analyzing genomic data as the amount of data has grown larger and more complex^8^. Applications of deep learning include gene annotation^9^, sequence pattern identification^10^, discovering biomarkers^11^, and inferring cell function and structure^12^. At its core, machine learning, and in particular deep learning, is utilized for classification based on training data and learning hidden patterns and structure in the data^13^. Although deep learning based approaches have been at the core of tremendous successes and popularity in many areas, including computer vision^14^, natural language processing^15^, and robotics^16^, it has some intrinsic disadvantages. First, it has limited explainability; models are often unable to fully detail the features and decision making process that led to a given classification or prediction^17^. Second, the computational cost and carbon footprint of such methods are skyrocketing while processing ever-increasing amounts of biological data^18^. Third, the predictions heavily rely on representative training data and the optimizations of hyperparameters^19^.

To address this, we introduce a fully transparent, efficient, explainable approach to assist end users in identifying the lab-of-origin of engineered DNA sequences. We solve the synthetic sequence tracking problem from an alignment perspective. Specifically, we make predictions by integrating the information of common sequences and “signature” sequences used in plasmid constructs via a pan-genome data structure. Pan-genomes have been studied for nearly two decades^20^. Pan genomes serve as a high-level summary of a biologically related group of genomes by capturing all of the group’s core and accessory regions, though the exact definition of a pan-genome can vary based on the application. In this paper, we define a pan-genome as a collection of genomic regions that are common or unique to synthetic biology research labs. Pan-genomes are frequently applied to capture the genomic diversity of a bacterial clade^21^. Many bioinformatic tools have been developed to build pan-genomes, such as Roary^22^, seq-seq-pan^23^, VG^24^, and *Plaster*^25^. To the best of our knowledge, *Plaster* is the only existing method that offers a linear time construction algorithm enabling it to scale to massive DNA sequence repositories. Building off of our prior work, we have developed a pan-genome for all available synthetic plasmid sequences using *Plaster*. We then use this synthetic sequence pan-genome as a framework for predicting the lab-of-origin of previously unseen, newly engineered sequences.

In this study, we demonstrate that pan-genome alignment combined with a lab score correction technique can successfully predict the lab-of-origin of an engineered DNA sequence 63% of the time. Around 80% of the time the source lab is included in the top 10 predicted labs. This approach has very few pre-processing steps, a quick update time for adding new known sequences, and a detailed and interpretable explanation for its predictions. This is in stark contrast to the previous Convolutional Neural Network (CNN) model which must be retrained to incorporate a single newly labeled sequence, and which is intrinsically a black box model.

## Results

### Neural Network vs *PlasmidHawk* Performance

We have developed a software called *PlasmidHawk* to predict the lab-of-origin of unknown synthetic DNA sequences. Lab-of- origin prediction with *PlasmidHawk* consists of three steps. The first step is to build a pan-genome from the synthetic plasmid training data using *Plaster*^25^. Second, in addition to building the pan-genome, *PlasmidHawk* annotates the pan-genome with records of which unique plasmid regions originated from which depositing labs. Lastly, *PlasmidHawk* predicts the lab-of-origin of new, unseen plasmids in the test data set by aligning those plasmids to the previously constructed and annotated pan-genome (Fig. 1).

**Figure 1.**
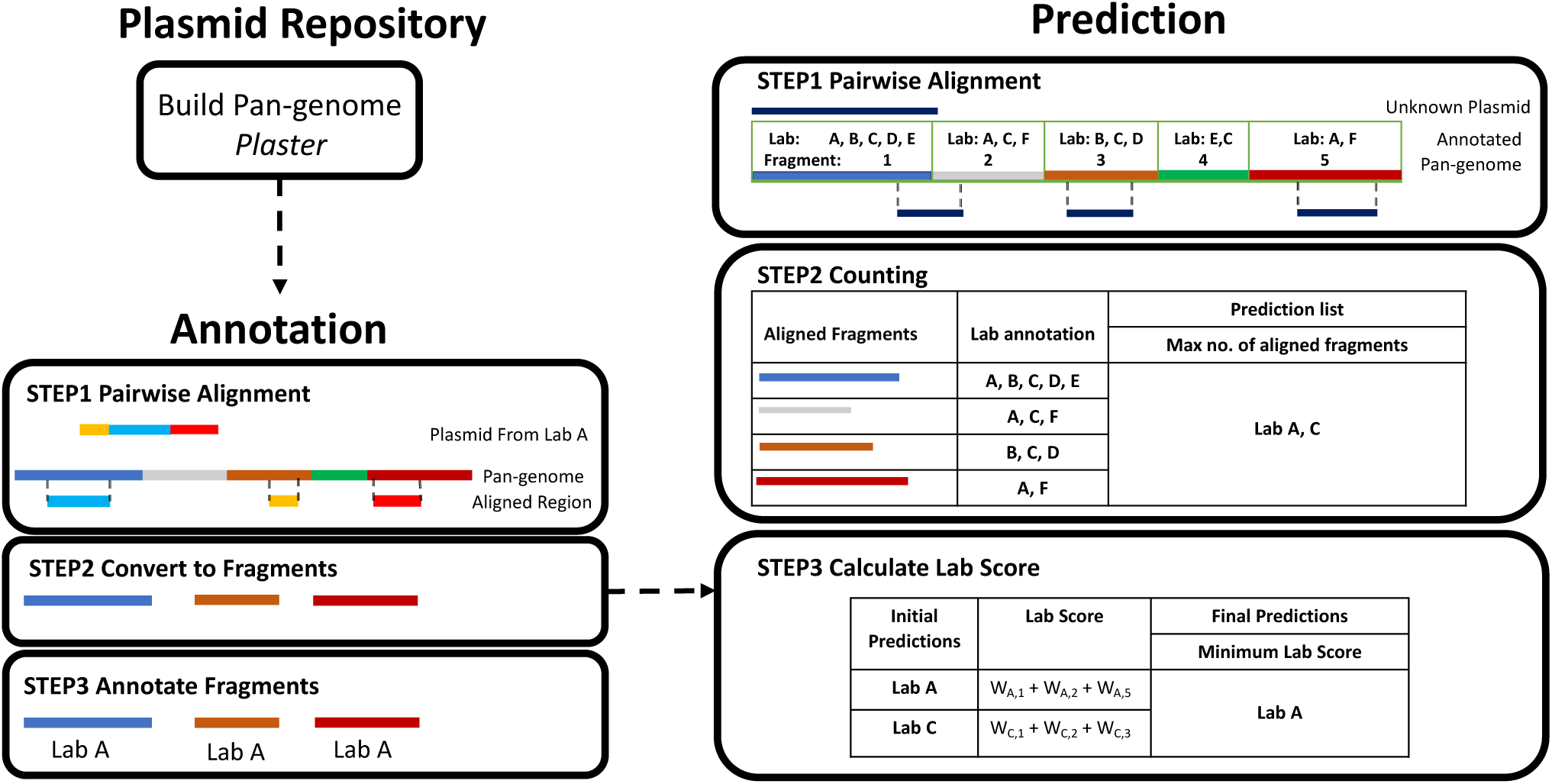
*PlasmidHawk* Pipeline: First, a pan-genome from the Addgene plasmids is built by *Plaster*. The plasmids are then aligned back to the final pan-genome to annotate the pan-genome with the depositing lab information for each aligned fragment. To predict the lab-of-origin of an unknown plasmid, *PlasmidHawk* aligns the unknown plasmid to the annotated pan-genome (Prediction Step 1) and counts the number of aligned fragments for each lab (Prediction Step 2). *PlasmidHawk* returns the labs that have the maximum number of aligned fragments as the predicted labs for lab-of-origin (Prediction Step 2). Finally, *PlasmidHawk* calculates the plasmid corrected score for each lab. This step takes the labs with the minimum corrected score as a final refinement step of its predictions for lab-of-origin (Prediction Step 3).

To begin the experiments, we select full plasmid sequences from labs who have at least ten deposited plasmids in Addgene for use as our input dataset. We split the sequences into two groups: the training group and the testing group. We use the training sequences to construct and annotate the synthetic plasmid pan-genome sequence *P*_*train*_. The pan-genome *P*_*train*_ contains a set of unique sequence fragments. Each fragment is denoted by its start and end positions in the pan-genome. Fragments are appended to one another creating a linear pan-genome with a delimiter sequence of N’s between consecutive fragments. Each fragment is further annotated with a list of the depositing labs who have sequences that align to the fragment.

After building and annotating the pan-genome, we predict the lab-of-origin of the sequences in the test set. In order to identify the lab-of-origin of an unknown plasmid, *PlasmidHawk* aligns a given test plasmid *p* to the input pan-genome and identifies the best aligned regions in the pan-genome (Fig. 1 Prediction Step 1). It then selects the pan-genome fragments that overlap with those aligned regions. We refer to these selected fragments as aligned fragments for the plasmid *p*. After identifying the aligned fragments, *PlasmidHawk* uses the fragments to predict the depositing lab or labs. Though *PlasmidHawk*, for instance in *MAXmode*, can return multiple labs as the most likely labs of origin, for this study we only allow one lab to be the true depositing lab. *PlasmidHawk* has two prediction modes: *MAXmode* and *CORRECTmode. MAXmode* predicts the lab-of-origin based on the set of labs who have the maximum number of aligned fragments for the plasmid *p* (Fig. 1 Prediction Step 2). Alternatively, *CORRECTmode* returns the lab or labs with the minimum “lab score”. *CORRECTmode* attempts to refine the set of labs from *MAXmode* by calculating the lab score which we introduce in this work. The lab score is calculated by weighting aligned fragments for each lab (Fig. 1 Prediction Step 3) (Methods).

To evaluate the performance of *PlasmidHawk*, we reran the deep learning experiments based on the description in Nielsen *et al*.^7^. We used the same data set used for the *PlasmidHawk* lab-of-origin prediction experiments. We construct and train the same CNN to predict the lab-of-origin of synthetic sequences. The final trained CNN can predict the correct lab-of-origin 36.5% of the time. The correct lab is ranked within the top 10 predicted labs 66.8% of the time (Fig. 2a). Our CNN prediction results are only slightly lower than the reported results in Nielsen *et al*., in which the accuracy is 48%, and 70% of the time the depositing lab is included in the top 10 predicted labs. Therefore, we believe our CNN experiments can fairly represent the performance of CNNs in lab-of-origin predictions. We believe the cause for the slight drop in performance between our CNN and the CNN built in Nielsen *et al*. is the larger size of our more up-to-date training dataset and the minor difference in how the true depositing source labs for each sequence were determined (Methods). We can further optimize the neural network to improve the results, but, in general, we do not expect a significant boost in prediction accuracy.

**Figure 2.**
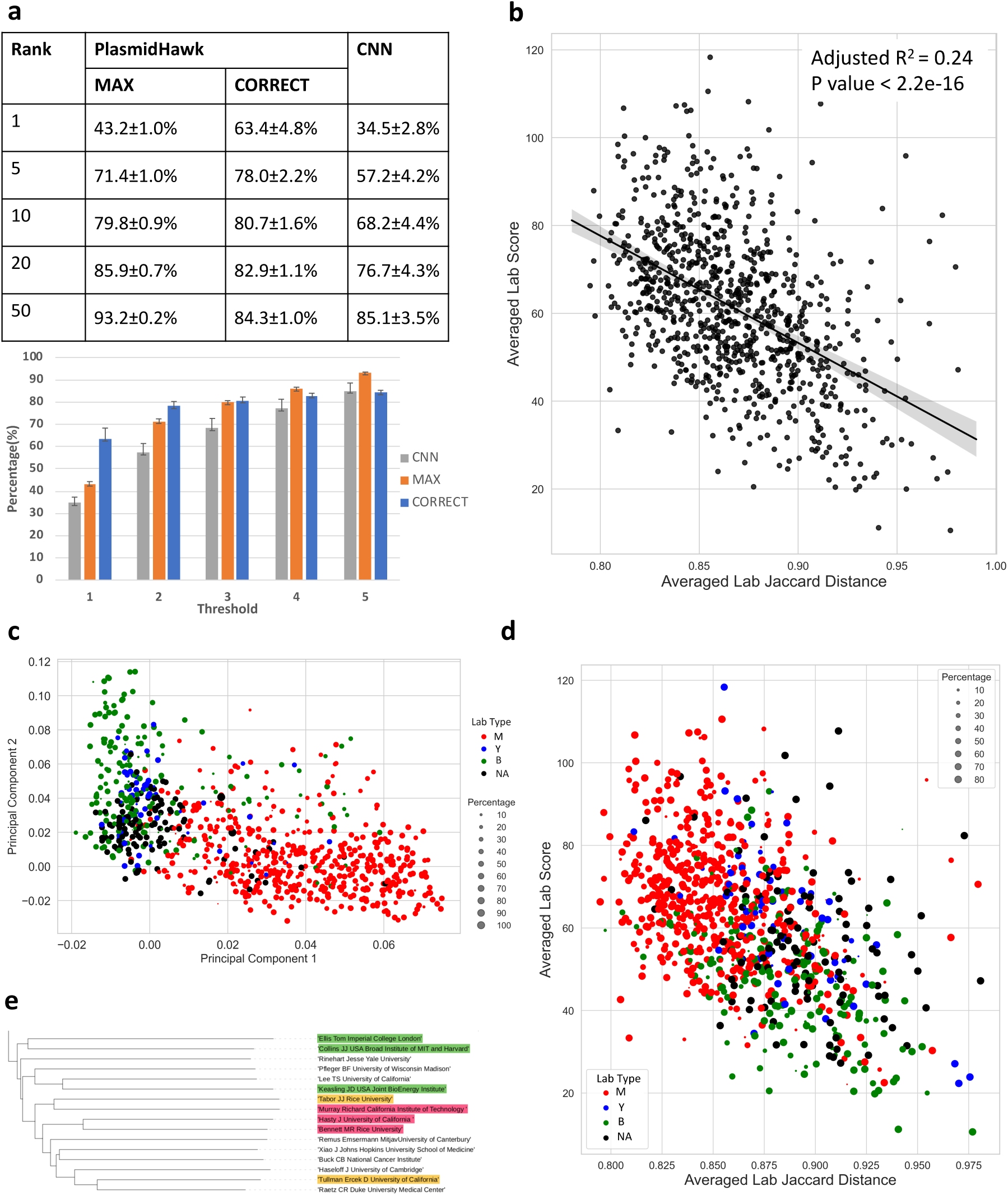
Prediction results and statistical analysis. **a** The performances of plasmid lab-of-origin prediction using *PlasmidHawk* and CNN. **b** Linear regression analysis between averaged lab Jaccard distances and averaged lab scores. Each dot represents a lab. The x axis shows averaged lab Jaccard distances. The larger the averaged lab Jaccard distance is, the more unique a lab’s plasmids are. The y axis is the averaged lab score. Labs with smaller averaged lab scores are more likely to be returned by *PlasmidHawk CORRECTmode* as predicted source labs. **c** Principal component analysis of labs based on lab Jaccard distances. The colors label labs based on their sequences’ host cells. The size of the dot corresponds to the percentage of the most abundant host cells inside a lab. Red: mammalian lab (M), blue: yeast lab (Y), green: bacterial lab (B), black: N/A lab (NA). **d** The distribution of different lab types visualized over the averaged lab Jaccard distance and averaged lab score for each lab. **e** A clade of the lab phylogenetic tree. Labs who belong to the same academic family or have collaborated with each other are highlighted with the same color.

### *MAXmode* ***accuracy***

To begin comparing the lab-of-origin prediction accuracy between the CNN and *PlasmidHawk MAXmode*, we only consider the *PlasmidHawk MAXmode* prediction results, considering only the case where the true source lab is reported as the single best prediction. A prediction result containing more than just the source lab in the highest scoring prediction set, or that does not contain the true source lab in this set, is classified as an incorrect prediction. In this case, *PlasmidHawk MAXmode* can reach 43.2% accuracy. To further compare the results between the CNN and *PlasmidHawk MAXmode*, we calculate the accuracies when considering the 5, 10, 20, and 50 best scoring predicted labs (Fig. 2a) (Methods). Overall, *PlasmidHawk MAXmode* behaves slightly better than the CNN does in all categories.

### *CORRECTmode* ***accuracy***

*PlasmidHawk CORRECTmode* identifies source labs by calculating lab scores for all the labs selected from *MAXmode* (Methods). The lab(s) with the lowest lab score values are considered as the final lab-of-origin predictions. Using the same test data sets used in the *MAXmode* prediction experiments, *PlasmidHawk CORRECTmode* has around 63% accuracy. 80% of the time the source lab is in the top 10 predicted labs (Fig. 2a). In addition, when compared to the CNN approach, the accuracies of both *MAXmode* and *CORRECTmode*, except for *CORRECTmode* with single lab predictions, have lower standard deviations (Fig. 2a). This means that in general lab-of-origin predictions from *PlasmidHawk* are more consistent.

### Lab Scores and Lab Sequence Similarities

*PlasmidHawk CORRECTmode* lab-of-origin experiments show that calculating the lab score can significantly improve the accurracies of predicting source labs (Fig. 2a). Lab scores are calculated by summarizing all the weights of different parts of query sequences. The magnitude of the weights are decided by the uniqueness of the sequences (Methods). *PlasmidHawk* then normalizes the lab scores and chooses labs with the minimum lab score as their final lab-of-origin prediction. Based on the workflow of *CORRECTmode*, we posit that labs with more unique sequences are more likely to have lower lab scores, that is, be identified as depositing labs.

To validate our hypothesis, we propose to describe the relationship between lab scores and the uniqueness of each lab using a regression model. First, we need to quantify the dissimilarities of a lab’s plasmids among other labs. To do that, we employ Jaccard distances, also called lab Jaccard distances in this paper, to estimate the differences between two labs’ sequences (Methods). A large pairwise lab Jaccard distance indicates there are few shared sequences between two labs. To summarize the idiosyncrasies of a lab’s sequences, we average all the pairwise lab Jaccard distances for individual labs, in order to generate a single value, referred as the averaged lab Jaccard distance, to represent the uniqueness of a lab’s sequences compared to the other 895 labs’ sequences. Additionally, we calculate averaged lab scores for each source lab (Methods). In general, the smaller averaged lab score a lab has, the more likely the lab is nominated as the true source lab by *PlasmidHawk CORRECTmode*.

After getting averaged lab scores and averaged Jaccard distances for all the depositing labs, We fit a linear regression model between these two variables. As averaged lab Jaccard distances increase, averaged lab scores decrease (P value *<* 2.2*e*− 16, adj*R*^2^ = 0.23) (Fig. 2b). The result shows that lab scores reflect the distinctness of labs sequences. It also indicates that *CORRECTmode* tends to link query sequences with labs who have more unique sequences.

### Lab Clustering and Lab Phylogenetic Tree

DNA engineering techniques have revolutionized the way people study biology. Among them, engineered plasmids are wildly used as vehicles to deliver and express non-native DNA into different host cells. At the beginning, scientists focused on expressing engineered plasmids inside bacteria. Recently, as DNA manipulation techniques have matured, people have shifted their attention from bacterial to mammalian cells in order to control and understand more complex cellular systems. Despite the rapid development in synthetic biology, there is no suitable approach to compare sequence similarities across different labs. The averaged lab Jaccard distance provides an initial attempt in comparing the uniqueness of individual labs’ sequences. In addition, the averaged lab Jaccard distance provides a way to quantify the development of plasmid engineering. Since one of the major trends in plasmid engineering is the shift from bacterial to mammalian hosts, in this section we evaluate the uniqueness of a lab’s sequences from a host cells perspective.

To do that, we first classify labs into different groups based on the highest percentage of their plasmids’ host cells. To simplify this problem, even though there are many different types of vector hosts shown in Addgene, we simply classify labs into four categories: mammalian (M), bacterial (B), yeast (Y) or N/A (NA). If a lab has more mammalian vectors than other types of vectors, the lab will be labelled as a mammalian lab (M). If a lab does not have a single plasmid belonging to bacterial, yeast or mammalian vectors, it is classified as N/A.

To identify the distribution of different target host cells across labs, we label their lab type in Fig. 2b to generate Fig. 2d. The size of each dot corresponds to the percentage of the prominent vector hosts for each lab (ties are resolved arbitrarily using lexicographic ordering of the types). For example, a lab classified as a mammalian lab could have only 20% mammalian vectors, with bacterial and yeast plasmids occupying less than or equal to 20% of the lab’s plasmids, and with the rest not belonging to any of the three host types we are considering (NA). Fig. 2d shows that labs focusing on engineering mammalian vectors are likely to have lower averaged lab Jaccard distances and higher averaged lab scores. Based on this result, we may roughly conclude that overall synthetic plasmids expressed in mammalian cells have lower sequence diversities and are less prone to have their lab-of-origin be identified by *PlasmidHawk CORRECTmode* than plasmids designed for yeast and bacterial cells.

In addition, we generated a PCA plot using the distance matrix of lab Jaccard distances for all labs and color the labs based on their host classification (Fig. 2c). The PCA plot shows a clear separation between mammalian labs and other types of labs along PC1. PC1 recovers the variation of host vector lab types and reveals these lab types as distinct, visually apparent clusters spanning across PC1. PC2 captures the variation within lab type group clusters. The principal component analysis further verifies our findings in Fig. 2d.

Furthermore, we construct a lab phylogenetic tree using lab Jaccard distances to reveal the academic relationships among all the labs (Methods). Figure 2e displays one of the clades of the lab phylogenetic tree. Branch lengths represent the distances between labs. In Figure 2e, principal investigators who belong to the same academic family or have collaborated with each other are highlighted by the same color. This indicates that the lab phylogenetic tree, which is derived from the alignment between the synthetic plasmid pan-genome and the original plasmid sequences, has the potential to reveal the academic genealogies in addition to being used for bioforensics.

### Comparisons with BLAST-based and CNN-based approaches

In Nielsen *et al*., researchers hand selected a plasmid from the Voigt lab (pCI-YFP,JQ394803.1) which exists in Genbank but not in the Addgene dataset, to compare the performances of the CNN and BLAST in identifying the source lab for an unknown sequence. BLAST fails to predict the lab-of-origin of pCI-YFP, ranking the Voigt lab 5th in its prediction. On the other hand, the CNN approach correctly predicted the plasmid with a significant p-value.

To evaluate *PlasmidHawk*’s performance, we predict the source lab for pCI-YFP using *PlasmidHawk*. We input the complete pan-genome *P*_*c*_, which compacts all the plasmids from labs who have at least 10 deposited full sequences in Addgene. Three labs, including Drew Endy, Ellington Andrew and Christopher Voigt, are identified as the set of top lab predictions using *PlasmidHawk MAXmode* (Fig.3a). It is worth noting here that this means that all three of these labs have the exact same amount of aligned fragments in the complete pan-genome *P*_*c*_.

**Figure 3.**
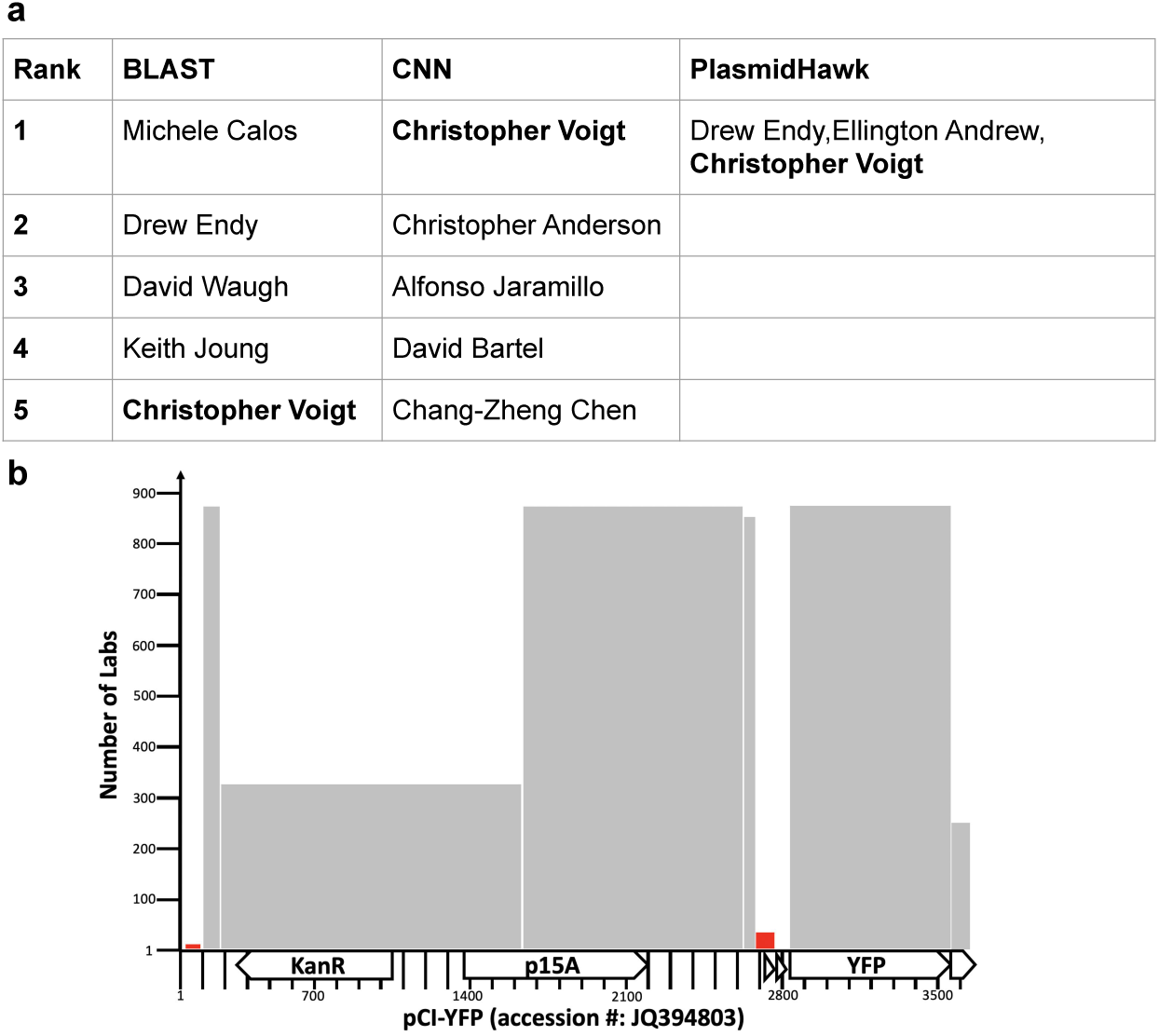
BLAST, CNN and *PlasmidHawk* prediction comparisons and interpretation **a** BLAST, CNN and *PlasmidHawk* lab-of-origin prediction results. **b** The number of labs annotated for the regions of pCI-YFP that aligned to different fragments in the synthetic pan-genome. Less than 20 labs have sequences aligned at 17-100nt and 2681-2750nt positions (red bars).

One of the advantages of using *PlasmidHawk* is that it can provide the alignment details of a given plasmid to explain the reasoning behind its prediction (Fig.3b). pCI-YFP aligns to a total of eight fragments in *P*_*c*_. The three labs selected by *PlasmidHawk* have all eight fragments. Among those eight fragments, six fragments have been used by more than 200 labs (grey bars in Fig. 3b). Those are common sequences used in the plasmid construction process. 17-100nt and 2681-2750nt in pCI-YFP are two regions that help *PlasmidHawk* narrow down the number of lab candidates (red bars in Fig. 3b). Only four labs have the fragment that aligned to 17-100nt in pCI-YFP. These labs are Drew Endy, Ellington Andrew, Christopher Voigt and Jaramillo Alfonso. 18 plasmids from Chirstopher Voigt, 2 plasmids from Ellington Andrew and 1 plasmids from Drew Endy in the training set align to the fragment (Supplementary Note Fig. 2). The 2 plasmids from Ellington Andrew were first published in Tabor *et al*.^26^, which Ellington Andrew and Christopher Voigt are co-authors of. In other words, 20 out of 21 training plasmids from the final predicted labs are from the Voigt lab.

In *PlasmidHawk CORRECTmode*, Drew Endy is the first prediction being returned for the lab-of-origin. Because the Voigt lab has the largest number of fragments and Drew Endy has the smallest number of fragments in the pan-genome among the three predicted labs, the lab score of the Voigt lab is the highest and the lab score of the Drew Endy lab is the lowest. Since the Voigt lab has the fourth most number of different fragments in the pan-genome, if there is more than one lab identified in *PlasmidHawk MAXmode*, the Voigt lab will typically rank lower in *PlasmidHawk CORRECTmode*. Based on the *PlasmidHawk CORRECTmode* lab-of-origin prediction experiments described above, 12 out of 15 times Voigt lab’s plasmids can be correctly identified. In all those 12 correct predictions, the Voigt lab is the only lab who has the maximum number of aligned fragments.

In general, the lab score can rank the source lab higher than the other labs identified in the *PlasmidHawk MAXmode*, but it is also biased towards picking labs with fewer fragments in the pan-genome. *PlasmidHawk* can return the predicted labs from *PlasmidHawk MAXmode* as well as their lab score for a given plasmid. Users have the ability to select the predicted labs based on their applications.

In the end, *PlasmidHawk* successfully predicted the correct lab for pCI-YFP as being in the set of labs containing the highest number of aligned fragments using *PlasmidHawk MAXmode*. In this case, *PlasmidHawk* has higher accuracy when compared to BLAST, but lower specificity when compared to the CNN. However, we again stress that *PlasmidHawk* additionally reveals its detailed decision making process and successfully identifies the signature sequences of the source lab. This human interpretable decision making process not only yields state of the art performance when analyzing all test set plasmids but also reveals the hidden intricacies of the pCI-YFP classification process. In an actual bioforensics setting, this type of evidence can be both crucial to a correct lab-of-origin assignment as well as necessary for a proper final justification which is human interpretable.

## Discussion

This study demonstrates that aligning unknown sequences to the synthetic plasmid pan-genome can effectively identify engineered DNA sequences’ lab-of-origin. Approaches based around knowledge of the problem coupled with standard tools such as simple alignment based methods can reach similar or better accuracy as state-of-the-art neural network based approaches. At the same time, *PlasmidHawk* provides deeply interpretable prediction results by providing unique sequence fragments responsible for identifying the source labs. As shown in the case of pCI-YFP, *PlasmidHawk* can even help elucidate a more in depth story of shared sequence fragments that are shared by many labs when constructing their synthetic plasmids as opposed to more unique regions used by very few labs. However, due to challenges such as these sequences that are commonly shared by many labs, about 60% of the time an unknown plasmid still cannot be successfully narrowed down to only the single correct depositing lab by *PlasmidHawk MAXmode*. To help ameliorate this difficulty, we have introduced the lab score for inferring the single correct lab-of-origin. The lab score helps order the set of predicted labs from *PlasmidHawk MAXmode* based on the pan-genome annotation. The incorporation of lab score increases the prediction accuracy from *MAXmode*’s 43.2% to *CORRECTmode*’s 63%.

Our work demonstrates an alignment-based approach can achieve high prediction accuracy while simultaneously providing detailed explanatory capabilities. Meanwhile, we are aware it has potential shortcomings versus a machine learning based approach. One issue with our method, for instance, is in determining the size and similarity thresholds for creating and adding novel fragments to the synthetic plasmid pan-genome. In a CNN model, more subtle signatures involving only one or a small handful of SNPs used by a particular lab can be captured, whereas this could potentially be overlooked by *PlasmidHawk*. Along these lines, plasmid design seems to be becoming more standardized^27^. In this standardization, sequences from a plasmid can be organized into different modules based on their functions. Researchers can then easily combine different modules to build a variety of plasmids. While this can significantly increase plasmid construction efficiency for synthetic biologists, it can have the unintended consequence of potentially weakening the available signal for determining the lab-of-origin, which may increase the concern of *PlasmidHawk*’s system using too coarse grained shared fragments as its primary source of signal for predictions.

Despite these concerns, along with the high prediction accuracy, *PlasmidHawk* preserves three main advantages over the deep learning approach for lab-of-origin prediction. First, the pan-genome alignment method has the ability to handle large amounts of input synthetic sequences quickly. Whenever a newly engineered sequence is created, the established synthetic plasmid pan-genome can be quickly updated by adding the unique parts of new sequences to the end of the pan-genome. The update time is less than 1s per plasmid^25^. On the other hand, for the deep learning approach, the CNN model has to be entirely re-trained whenever there are new sequences coming in. As more and more synthetic sequences are being added into the database, the computational and environmental cost for neural network approaches will increase alongside^18^. Second, *PlasmidHawk* is not restricted to predict labs with a small number of deposited plasmids. It has the potential to identify a given plasmid as having originated from a lab that only has one recorded plasmid in the training data set. On the other hand, the CNN requires labs to have enough plasmids in the training set to then be able to be predicted. Third, the pan-genome alignment approach can associate specific regions of an unknown sequence to the source lab. By doing this, *PlasmidHawk* provides detailed evidence to support its lab classifications. When combined, these three benefits create a white box approach that, when compared to the deep learning method, make *PlasmidHawk* a welcome addition to the current methods in biodefense and bioforensics.

Finally, we have demonstrated a new way to characterize research diversity and relationships among synthetic biology labs. By aligning synthetic plasmids from each depositing lab to the synthetic plasmid pan-genome, we are able to capture the resemblances and variations between all labs engaged in synthetic plasmid engineering. The lab phylogenetic tree we have created not only reveals the research structure in synthetic biology, but also implies academic lineage and collaborations. More work on comparative genomics approaches based on phylogenetics can further assist to trace back the lab-of-origin of an unknown plasmid.

Overall, the aim of our study is not to diminish the achievement of deep learning in genomic analyses, but to point out the value of investigating traditional comparative genomics methods such as by using genome alignment. We believe a truly optimal method would combine the benefits of both well studied and interpretable methods like genome alignment with the power of deep learning models such as the CNN model of Nielsen *et al*..

## Methods

### Addgene Dataset

We acquired a plasmid dataset from Addgene in January 2019. Addgene is a synthetic plasmid repository. It was used in Nielsen *et al*. to conduct the deep learning lab-of-origin prediction study. DNA sequences in Addgene can be classified into four categories: full plasmid sequences submitted by Addgene, partial sequences submitted by Addgene, full plasmid sequences submitted by a depositing lab, and partial sequences submitted by a depositing lab. There are a total 51, 047 complete sequences, in which 28, 879 are uploaded by Addgene, and 73,7272 partial sequences. The DNA sequences and their metadata are stored in a JavaScript Object Notation (JSON) file. In Nielsen *et al*., a plasmid depositing lab was parsed directly from the JSON file. However, the JSON file we obtained had no deposting lab information. To decide a plasmid’s depositing lab, we first found information from its Genbank file. We took the last author in the first published paper of the plasmid as the depositing lab. For the plasmids without Genbank files, we looked up the author information through its PubMed ID (PMID) or PMCID in the JSON file. If we still could not find its depositing lab, we parsed the depositor information directly from the Addgene website.

### *PlasmidHawk* **Workflow**

The main goal of our study is to predict engineered plasmids’ lab-of-origin by aligning unknown plasmids to a synthetic plasmid pan-genome. In order to do that, we developed a lab-of-origin prediction software called *PlasmidHawk*. It consists of three modules: pan-genome construction, pan-genome annotation, and lab-of-origin prediction. In general, the three modules should be used sequentially for plasmid lab-of-origin detection. Each module can also be applied independently for other scientific purposes.

For the first module, *PlasmidHawk* takes in plasmids from the Addgene database and constructs a synthetic plasmid pan-genome *P* using *Plaster*. *Plaster* is a state-of-the-art linear pan-genome construction algorithm^25^. The final pan-genome *P* is composed of a set of sequence fragments *F* = [*f*_0_, *f*_1_, … *f*_*n*_]. Each fragment is at least 50bp long (default) and connected to neighboring fragments by a delimiter sequence of 10 “N” nucleotides. After building the synthetic plasmid pan-genome, *PlasmidHawk* aligns input plasmids back to the pan-genome. Plasmid sequences extending over a defined fragment are separated. If a pan-genome fragment has at least 20bp (default) matching with sequences from an input plasmid, the fragment is annotated with that plasmid’s depositing lab.

To predict the lab-of-origin of an unknown plasmid *p, PlasmidHawk* aligns the plasmid *p* to the reference pan-genome *P* built in the first step. It then extracts aligned pan-genome fragments from the pan-genome. Each aligned pan-genome fragment has a match of at least 20bp with the plasmid *p. PlasmidHawk MAXmode* then outputs lab(s) with the highest number of aligned fragments as the predicted source lab(s). To further improve the lab-of-origin prediction accuracy, *PlasmidHawk CORRECTmode* calculates lab scores for labs returned from *PlasmidHawk MAXmode. CORRECTmode* takes lab(s) with the minimum lab score as the final prediction as explained in full detail in the following section and visualized in Fig. 4. Briefly, the value of a lab’s score depends on the total number of pan-genome fragments each lab has and the number of labs sharing the same aligned fragments. Mathematically, the lab score for lab *l*, denoted as *S*_*l*_, is

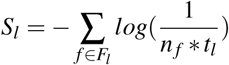

where *F*_*l*_ is the aligned fragments set for lab *l*. It includes all the aligned fragments lab *l* has for the query sequence. *n*_*f*_ is the number of labs sharing fragment *f* in the pan-genome *P*. And *t*_*l*_ is the total number of fragments lab *l* has in *P*.

**Figure 4.**
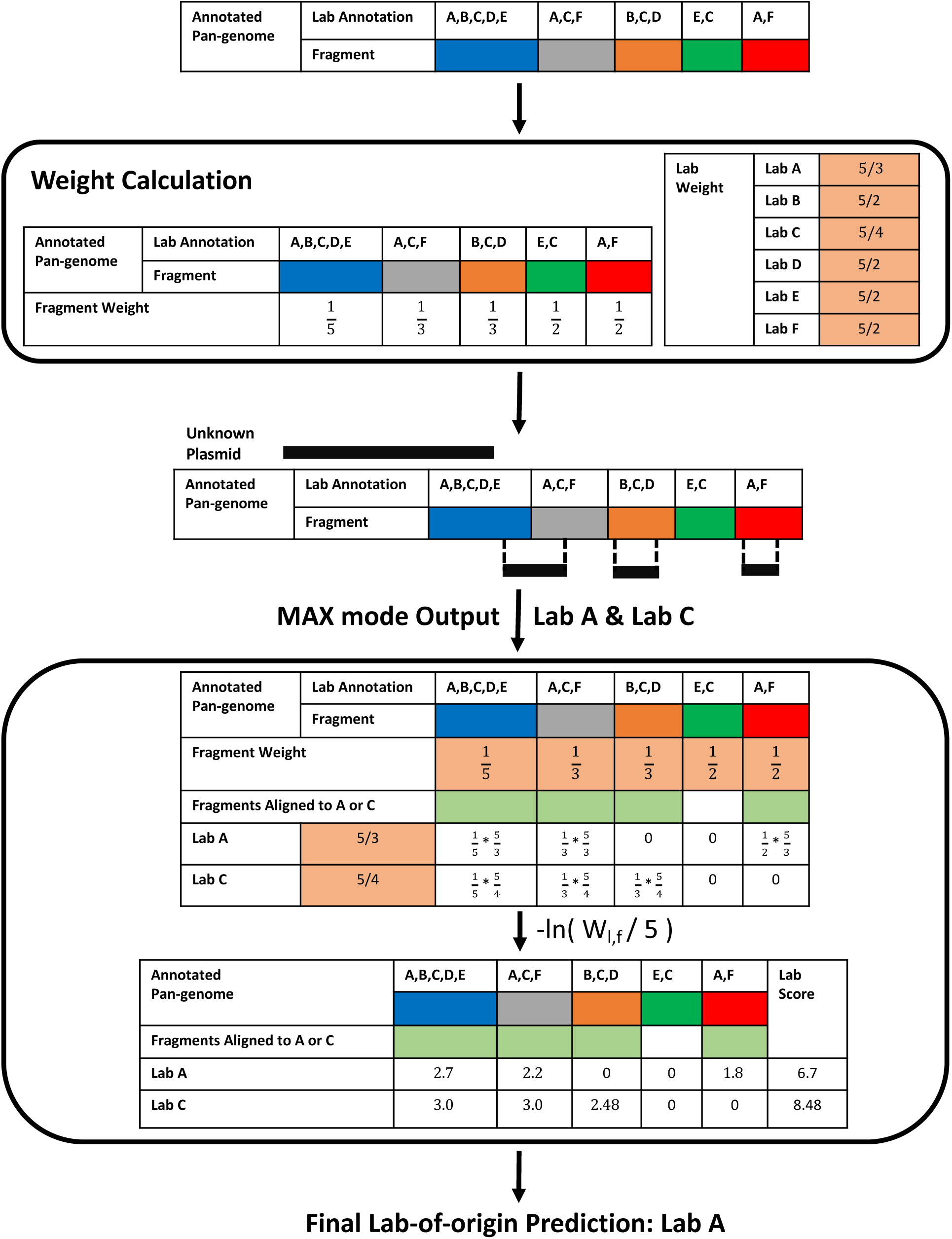
*PlasmidHawk CORRECTmode* workflow. After building the reference pan-genome, *CORRECTmode* calculates fragment weights and lab weights according to the pan-genome annotations. To predict the lab-of-origin of an unknown sequence, *PlasmidHawk* first aligns the query sequence to the pan-genome and identifies candidate source labs through *MAXmode*: in this case, lab A and lab C. *CORRECTmode* then calculates lab scores for the labs output from *MAXmode*. In the end, *CORRECTmode* predicts the lab with the minimum lab score, lab A, as the depositing lab.

### Lab Score

After obtaining a list of potential source labs from *MAXmode, PlasmidHawk CORRECTmode* attempts to further narrow down the labs to the true source lab. To do that, it calculates lab scores to rank labs returned from *MAXmode* as shown in Fig. 4. Essentially, lab scores are assigned to individual labs through a weighting function. Labs with lower lab scores have a higher chance to be the real depositing labs. The weighting function used to calculate lab scores is derived from our key observations that despite the maximum number of aligned fragments being the same among multiple labs, pan-genome fragments shared by many labs are potentially less informative versus fragments shared by few labs. Also, labs with less total fragments in the pan-genome can be weighted higher when making the final predictions.

Specifically, after constructing and annotating the reference pan-genome, *CORRECTmode* first calculates weights for each lab, denoted as *W*_*l*_ for lab *l*, and each fragment, referred as *W*_*f*_ for fragment *f*, based on the pan-genome annotations. The lab weight 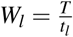, where *T* is the total number of fragments in the reference pan-genome and *t*_*l*_ is the total number of fragments lab *l* has in the pan-genome. *W*_*l*_ is the reciprocal of the fraction of fragments annotated by lab *l* inside the pan-genome. The fragment weight 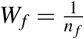, where *n* _*f*_ is the total number of labs annotated to fragment *f* (Fig. 4).

To calculate the final scores used for prediction, *CORRECTmode* goes through all the labs returned by *MAXmode* and all the aligned fragments for each of these labs and calculates a joint weight *W*_*l, f*_ for each aligned fragment *f* and lab *l*. To calculate *W*_*l, f*_, *PlasmidHawk* first identifies the set of aligned fragments *F*_*l*_ for each lab *l* identified by *MAXmode. CORRECTmode* then calculates *W*_*l, f*_ by multiplying *W*_*l*_ and *W*_*f*_, and normalizing it with its maximum possible value (*T*). The normalization bounds *W*_*l, f*_ between 0 and 1. Mathematically,

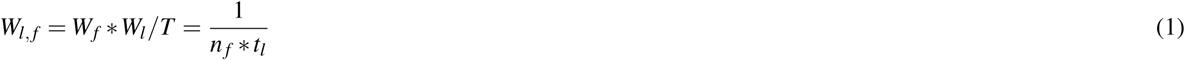

*CORRECTmode* then does a log transformation of each joint weight to avoid multiplication operations and potential overflows when calculating the final single fragment lab score *S*_*l, f*_. It adds a final negative sign to each transformed value to make the final scores positive.

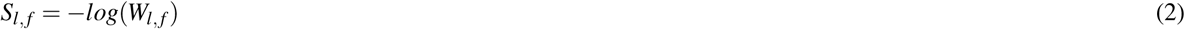

Finally, *CORRECTmode* sums the *S*_*l, f*_ of all the fragments in *F*_*l*_ to generate the final lab score *S*_*l*_ for lab *l* used for the final prediction. The lab with the lowest *S*_*l*_ is chosen as the predicted lab-of-origin as outlined in Fig. 4.

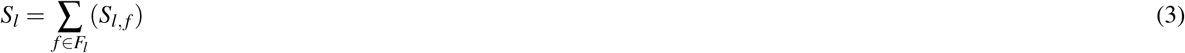

### *PlasmidHawk* **Lab of Origin Prediction**

Given the nature of neural networks, the CNN needs a certain number of plasmids from each lab to train, test and validate the neural network. Although *PlasmidHawk* does not have this kind of requirement, in order to have a fair comparison between the CNN and *PlasmidHawk*, we only choose labs with at least 10 complete plasmids (“Full Repository” or “Full Depositor”) sequences in Addgene to conduct this experiment. A total of 38, 681 plasmid sequences from 896 labs are used in this experiment. To evaluate the performance of *PlasmidHawk*, sequences are split into two groups: three plasmids from each lab are randomly selected for testing and the remaining plasmids are used for the training set. We utilize the training plasmids to build and annotate the synthetic plasmid pan-genome. We evaluate *PlasmidHawk* prediction accuracy using plasmids in the test data set. The entire process is repeated five times.

To assess the lab-of-origin prediction accuracies of *MAXmode* and *CORRECTmode*, we test *PlasmidHawk* at different thresholds; we test the accuracy when considering the top 1, 5, 10, 20, and 50 labs output from *PlasmidHawk*. For *MAXmode*, we only consider the top predictions at or above the threshold, and eliminate sets of predictions whose inclusion would cause there to be more predictions than the threshold. For *CORRECTmode*, we only consider the scored, ordered list created from the top set of predictions from *MAXmode*. For example, when setting the threshold at 1, for *MAXmode*, we only consider correct predictions when the top set of predicted labs contains only a single lab and that is the correct depositing lab. If *MAXmode* outputs more labs in the top set of predictions than the threshold, no labs are considered and the prediction is considered incorrect, even if the correct lab is in the top set of predictions. For *CORRECTmode* with a threshold of 1, we order the top set of *MAXmode* predictions, and only consider the single best scoring prediction. As another example, when setting the threshold at 5, and *MAXmode* outputs a set of two labs as the top predictions, two labs in the second best set of predictions, and two labs in the third best set of predictions, the four labs in the top two sets would be considered and the two labs in the third set would not be considered. In this and all other cases, *CORRECTmode* considers only the top set of labs from *MAXmode*, thus for the top 5 threshold it would still only consider the ranked list of the top two labs from *MAXmode*. In addition, if a set of labs in *CORRECTmode* have the same lab score and their rankings are around the threshold, we would arbitrarily select few labs from the set and add them into our final prediction lists. By doing this, the number of labs in a *CORRECTmode* prediction result equals to the threshold. For instance, if the threshold is 1 and there are two labs with the same lab score returned from *CORRECTmode*, we will arbitrarily select one lab as the *CORRECTmode* prediction.

### Convolutional Neural Network Architecture and Lab of Origin Prediction

The CNN architecture was constructed based on Nielsen *et al*. All the CNN parameters are set to their optimized values from Nielsen *et al*.. In the original experimental design, the authors reported splitting the data set into six subsets because of memory limitations. We replicate this by separating the training data into six subsets, then load and train one at a time in each epoch. After training, we save the final model and parameters.

We use the same plasmid data set from the *PlasmidHawk* experiments to train, validate and test the CNN approach. We randomly pick three plasmids from each lab as the validation data set and then pick additional three plasmids as the test data set. The remaining plasmids are used as the training set. We preprocess and encode the DNA sequences. As in Nielsen *et al*., we set all the DNA sequences to 8000bp long by truncating the long DNA sequences and padding the short DNA sequences with Ns. Characters other than A, T, C, or G in the DNA sequences are converted to Ns. We append the sequence’s reverse complement to itself with 48 Ns in between. We translate those processed sequences as a one-hot vector with length 16,048 where A=[1000], T=[0100], G=[0010],C=[0000]. The depositing lab is encoded as a one-hot vector with total length 896. This experiment is repeated five times.

To evaluate the CNN prediction accuracy, we calculate the percentage of plasmids in the test data set correctly identified their lab-of-origin while considering the top 1, 5, 10, 20, and 50 predicted labs from CNN. We then compute the average percentages of correct predictions and their standard deviations at different thresholds (1, 5, 10, 20, 50).

### Averaged Lab Jaccard Distance and Averaged Lab Score

To evaluate the relationships between lab scores and the uniqueness of labs’ plasmids, we calculate the lab Jaccard distance between all labs. The lab Jaccard distance quantifies the sequence similarities between all plasmids between two labs. To measure lab Jaccard distances, we first build and annotate a complete synthetic plasmid pan-genome using all plasmids from labs who have at least 10 complete sequences. We then extract all the fragments annotated with lab *A* to fragment set *F*_*A*_. We do this again for all labs. We define the lab Jaccard distance between two labs, lab *A* and lab *B*, as *JD*(*A, B*) = 1 −*J*(*F*_*A*_, *F*_*B*_), where 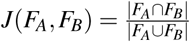 represents the Jaccard index between two labs (Fig.5). We built a distance matrix between all labs by calculating the pairwise Jaccard distances between every pair of labs. This distance matrix was used, for instance, to build the lab phylogenetic tree and also to calculate the “averaged lab Jaccard distance” for each individual lab. The averaged lab Jaccard distance for a lab is simply the average of all the cells in the corresponding row or column for that lab in the distance matrix.

**Figure 5.**
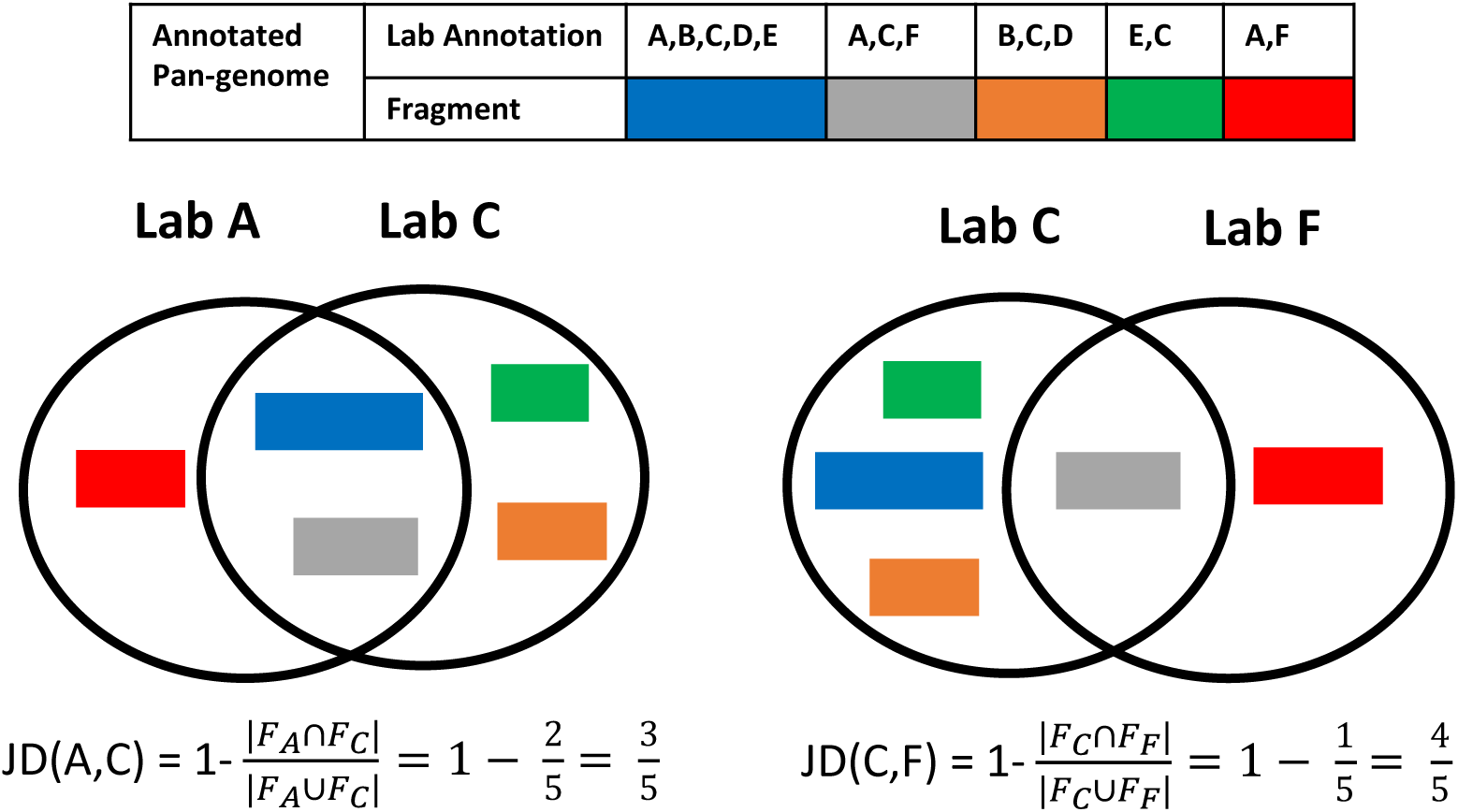
Lab Jaccard distance calculation. To calculate lab Jaccard distances between two labs, such as lab A and lab C, we first build a fragment set, *F*_*A*_ and *F*_*C*_, for each lab. A fragment set contains all the pan-genome fragments annotated by the corresponding labs. The lab Jaccard distance between lab *A* and lab *C* is *JD*(*A,C*) = 1 − *J*(*F*_*A*_, *F*_*C*_), where the Jaccard index (*J*(*F*_*A*_, *F*_*C*_)) is the fraction of shared pan-genome fragments out of all the different fragments lab A and C have. A large lab Jaccard distance represents two labs have few shared sequences.

To calculate a lab’s “averaged lab score”, we first run *CORRECTmode* on all test plasmids from the five independent lab-of-origin prediction experiments. If *CORRECTmode* returns a score for the true depositing lab for a plasmid, we assign that returned lab score to that plasmid. A lab’s “averaged lab score” is the average of all assigned lab scores for all test plasmids corresponding to that lab.

### pCI-YFP Prediction Analysis

To identify pCI-YFP’s depositing lab, we input the pCI-YFP sequence, the complete synthetic pan-genome sequence, and the pan-genome annotation information into *PlasmidHawk. PlasmidHawk* returns the predicted labs and their lab scores. It also outputs alignment results. We retrieve the aligned pan-genome fragments of pCI-YFP and the list of labs having those aligned fragments to create the alignment plot (Fig. 3b). In addition, we align all the plasmids that are used in building the complete pan-genome and have shared sequences with pCI-YFP from 17-100nt.

### Lab Phylogenetic Tree

We apply RapidNJ^28^ to construct a lab phylogenetic tree based on lab Jaccard distances. RapidNJ employs the neighbour-joining method^29^ to build phylogenetic trees. The visualization is conducted with the interactive tree of life (https://itol.embl.de)^30^. The full lab phylogenetic tree can be viewed in: http://itol.embl.de/shared/qiwangrice.

## Statistical Analysis

The principal component analysis is conducted using the sklearn.decomposition function^31^. The explained variances for PC1 and PC2 are 2.1 and 0.4. All the code is available in the gitlab repository.

## Code availability

PlasmidHawk is written in Python and is available at https://gitlab.com/treangenlab/plasmidhawk.git

## Data availability

All results supporting this study are available on Gitlab. The raw sequences used in this study are available upon request.

## Acknowledgements

Q. W. was supported by funds from Rice University and by funds from the National Institute for Neurological Disorders and Stroke (NINDS) of the National Institutes of Health under award number R21NS106640. T.R.L. was supported by funds from Rice University. R. A. L. E. was supported by the FunGCAT program from the Office of the Director of National Intelligence (ODNI), Intelligence Advanced Research Projects Activity (IARPA), via the Army Research Office (ARO) under Federal Award No. W911NF-17-2-0089. T.J.T was supported by startup funds from Rice University and was partially supported by the FunGCAT program from the Office of the Director of National Intelligence (ODNI), Intelligence Advanced Research Projects Activity (IARPA), via the Army Research Office (ARO) under Federal Award No. W911NF-17-2-0089.

The authors would like to thank Dr. Caleb Bashor and Dr. Tianzhou Ma for critical discussion and feedback, and Dr. Joanne Kamens from Addgene for providing prompt access to the DNA sequences of the synthetic plasmids utilized in this study. The views and conclusions contained herein are those of the authors and should not be interpreted as necessarily representing the official policies or endorsements, either expressed or implied, of the ODNI, IARPA, ARO, or the US Government.

## Author contributions statement

All authors conceived the experiment(s), analysed the results and reviewed the manuscript. Q.W. and T.R.L. conducted the experiment(s). T.J.T. managed the project.

The corresponding author T.J.T. is responsible for submitting a competing interests statement on behalf of all authors of the paper.

