## Supplementary File 1 for "PlasmidHawk: Alignment-based Lab-of-Origin Prediction of Synthetic Plasmids"

**Supplementary Note 1** Because synthetic biology labs have a wide range of sequence diversities and vector types, the percentage of correctly predicted plasmids out of all the test plasmids from a lab varies across labs. The percentage of correct predictions for each lab is referred as lab sensitivity. Lab sensitivity (all) counts all the plasmids correctly predicted by $PlasmidHawk$ regardless of the number of plasmids returned from $PlasmidHawk$. On the other hand, lab sensitivity (single) only takes $CORRECTmode$ predictions with single true source lab as right predictions. Since in each prediction experiment we randomly choose three plasmids from each lab to detect their lab-of-origin, the same plasmids can be taken as test plasmids in more than one experiments. Therefore, while calculating lab sensitivities, the same plasmids can be counted multiple times.

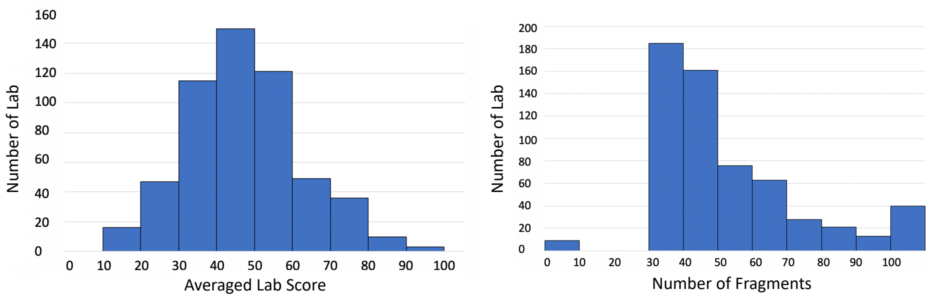


Figure 1: **a** The distribution of averaged lab scores. The median of the averaged lab score is 46. **b** The distribution of the number of pan-genome fragments each lab has. Most of the labs have from 30 to 50 number of fragments in the pan-genome.

| Group | Plasmid | Start | Sequence | End |
| --- | --- | --- | --- | --- |
| | pCI-YFP | 17 | TGCAGTCCGGCAAAAAAACGGGCAAGGTGTCACCACCCTGCCCTTTTTCTTTAAAACCGAAAAGATTACTTCGCGTTATGCAGG | 100 |
| Christopher Voigt | 60589_user | 1628 | TGCAGTCCGGCAAAAAAACGGGCAAGGTGTCACCACCCTGCCCTTTTTCTTTAAAACCGAAAAGATTACTTCGCGTTATGCAGG | 1545 |
| | | 3065 | TCCGGCAAAAAAACGGGCAAGGTGTCACCACCCTGCCCTTTTTCTTTAAAACCGAAAAGATTACTTCGCGTT-TGC | 3139 |
| | 60588_user | 1628 | TGCAGTCCGGCAAAAAAACGGGCAAGGTGTCACCACCCTGCCCTTTTTCTTTAAAACCGAAAAGATTACTTCGCGTTATGCAGG | 1545 |
| | | 3065 | TCCGGCAAAAAAACGGGCAAGGTGTCACCACCCTGCCCTTTTTCTTTAAAACCGAAAAGATTACTTCGCGTT-TGC | 3139 |
| | 60587_user | 1628 | TGCAGTCCGGCAAAAAAACGGGCAAGGTGTCACCACCCTGCCCTTTTTCTTTAAAACCGAAAAGATTACTTCGCGTTATGCAGG | 1545 |
| | | 3065 | TCCGGCAAAAAAACGGGCAAGGTGTCACCACCCTGCCCTTTTTCTTTAAAACCGAAAAGATTACTTCGCGTT-TGC | 3139 |
| | 60586_user | 1628 | TGCAGTCCGGCAAAAAAACGGGCAAGGTGTCACCACCCTGCCCTTTTTCTTTAAAACCGAAAAGATTACTTCGCGTTATGCAGG | 1545 |
| | | 3065 | TCCGGCAAAAAAACGGGCAAGGTGTCACCACCCTGCCCTTTTTCTTTAAAACCGAAAAGATTACTTCGCGTT-TGC | 3139 |
| | 60574_user | 1628 | TGCAGTCCGGCAAAAAAACGGGCAAGGTGTCACCACCCTGCCCTTTTTCTTTAAAACCGAAAAGATTACTTCGCGTTATGCAGG | 1545 |
| | | 3086 | TCCGGCAAAAAAACGGGCAAGGTGTCACCACCCTGCCCTTTTTCTTTAAAACCGAAAAGATTACTTCGCGTT-TGC | 3160 |
| | 60575_user | 1649 | TGCAGTCCGGCAAAAAAACGGGCAAGGTGTCACCACCCTGCCCTTTTTCTTTAAAACCGAAAAGATTACTTCGCGTTATGCAGG | 1566 |
| | | 3086 | TCCGGCAAAAAAACGGGCAAGGTGTCACCACCCTGCCCTTTTTCTTTAAAACCGAAAAGATTACTTCGCGTT-TGC | 3160 |
| | 60576_user | 1649 | TGCAGTCCGGCAAAAAAACGGGCAAGGTGTCACCACCCTGCCCTTTTTCTTTAAAACCGAAAAGATTACTTCGCGTTATGCAGG | 1566 |
| | | 3086 | TCCGGCAAAAAAACGGGCAAGGTGTCACCACCCTGCCCTTTTTCTTTAAAACCGAAAAGATTACTTCGCGTT-TGC | 3160 |
| | 60577_user | 1649 | TGCAGTCCGGCAAAAAAACGGGCAAGGTGTCACCACCCTGCCCTTTTTCTTTAAAACCGAAAAGATTACTTCGCGTTATGCAGG | 1566 |
| | | 3086 | TCCGGCAAAAAAACGGGCAAGGTGTCACCACCCTGCCCTTTTTCTTTAAAACCGAAAAGATTACTTCGCGTT-TGC | 3160 |
| | 60578_user | 1649 | TGCAGTCCGGCAAAAAAACGGGCAAGGTGTCACCACCCTGCCCTTTTTCTTTAAAACCGAAAAGATTACTTCGCGTTATGCAGG | 1566 |
| | | 3086 | TCCGGCAAAAAAACGGGCAAGGTGTCACCACCCTGCCCTTTTTCTTTAAAACCGAAAAGATTACTTCGCGTT-TGC | 3160 |
| | 60579_user | 1649 | TGCAGTCCGGCAAAAAAACGGGCAAGGTGTCACCACCCTGCCCTTTTTCTTTAAAACCGAAAAGATTACTTCGCGTTATGCAGG | 1566 |
| | | 3086 | TCCGGCAAAAAAACGGGCAAGGTGTCACCACCCTGCCCTTTTTCTTTAAAACCGAAAAGATTACTTCGCGTT-TGC | 3160 |
| | 49375_user | 1649 | TGCAGTCCGGCAAAAAAACGGGCAAGGTGTCACCACCCTGCCCTTTTTCTTTAAAACCGAAAAGATTACTTCGCGTTATGCAGG | 1566 |
| | 60581_user | 1649 | TGCAGTCCGGCAAAAAAACGGGCAAGGTGTCACCACCCTGCCCTTTTTCTTTAAAACCGAAAAGATTACTTCGCGTTATGCAGG | 1566 |
| | | 3086 | TCCGGCAAAAAAACGGGCAAGGTGTCACCACCCTGCCCTTTTTCTTTAAAACCGAAAAGATTACTTCGCGTT-TGC | 3160 |
| | 60580_user | 1649 | TGCAGTCCGGCAAAAAAACGGGCAAGGTGTCACCACCCTGCCCTTTTTCTTTAAAACCGAAAAGATTACTTCGCGTTATGCAGG | 1566 |
| | | 3086 | TCCGGCAAAAAAACGGGCAAGGTGTCACCACCCTGCCCTTTTTCTTTAAAACCGAAAAGATTACTTCGCGTT-TGC | 3160 |
| | 60583_user | 1649 | TGCAGTCCGGCAAAAAAACGGGCAAGGTGTCACCACCCTGCCCTTTTTCTTTAAAACCGAAAAGATTACTTCGCGTTATGCAGG | 1566 |
| | | 3086 | TCCGGCAAAAAAACGGGCAAGGTGTCACCACCCTGCCCTTTTTCTTTAAAACCGAAAAGATTACTTCGCGTT-TGC | 3160 |
| | 60582_user | 1649 | TGCAGTCCGGCAAAAAAACGGGCAAGGTGTCACCACCCTGCCCTTTTTCTTTAAAACCGAAAAGATTACTTCGCGTTATGCAGG | 1566 |
| | | 3086 | TCCGGCAAAAAAACGGGCAAGGTGTCACCACCCTGCCCTTTTTCTTTAAAACCGAAAAGATTACTTCGCGTT-TGC | 3160 |
| | 60584_user | 1649 | TGCAGTCCGGCAAAAAAACGGGCAAGGTGTCACCACCCTGCCCTTTTTCTTTAAAACCGAAAAGATTACTTCGCGTTATGCAGG | 1566 |
| | | 3086 | TCCGGCAAAAAAACGGGCAAGGTGTCACCACCCTGCCCTTTTTCTTTAAAACCGAAAAGATTACTTCGCGTT-TGC | 3160 |
| | 60594_user | 3584 | TGCAGTCCGGCAAAAAAACGGGCAAGGTGTCACCACCCTGCCCTTTTTCTTTAAAACCGAAAAGATTACTTCGCGTTATGCAGG | 3501 |
| | | 5021 | TCCGGCAAAAAAACGGGCAAGGTGTCACCACCCTGCCCTTTTTCTTTAAAACCGAAAAGATTACTTCGCGTT-TGC | 5095 |
| | 31392_user | 4407 | TGCAGTCCGGCAAAAAAACGGGCAAGGTGTCACCACCCTGCCCTTTTTCTTTAAAACCGAAAAGATTACTTCGCGTTATGCAGG | 4490 |
| Ellington Andrew | 25088_user | 5438 | TGCAGTCCGGCAAAAAAACGGGCAAGGTGTCACCACCCTGCCCTTTTTCTTTAAAACCGAAAAGATTACTTCGCGTTATGCAGG | 5521 |
| | 25092_user | 6082 | TGCAGTCCGGCAAAAAAACGGGCAAGGTGTCACCACCCTGCCCTTTTTCTTTAAAACCGAAAAGATTACTTCGCGTTATGCAGG | 6165 |
| Drew Endy | 44456_user | 4528 | TGCAGTCCGGCAAAAAAACGGGCAAGGTGTCACCACCCTGCCCTTTTTCTTTAAAACCGAAAAGATTACTTCGCGTTATGCAGG | 4611 |

Figure 2: Multiple sequence alignment of training plasmids and pC-YFP at 17-100nt. In the training data set, 18 plasmids from Chirstopher Voigt, 2 plasmids from Ellington Andrew and 1 plasmids from Drew Endy align to the fragment. The 2 plasmids from Ellington Andrew were engineered by collaborating with Christopher Voigt. For the 18 training plasmids from Voigt lab, each plasmid have two parts of sequences aligned to the pan-genome fragment. One part has the exact matches, while the other part has one nucleotide deletion.

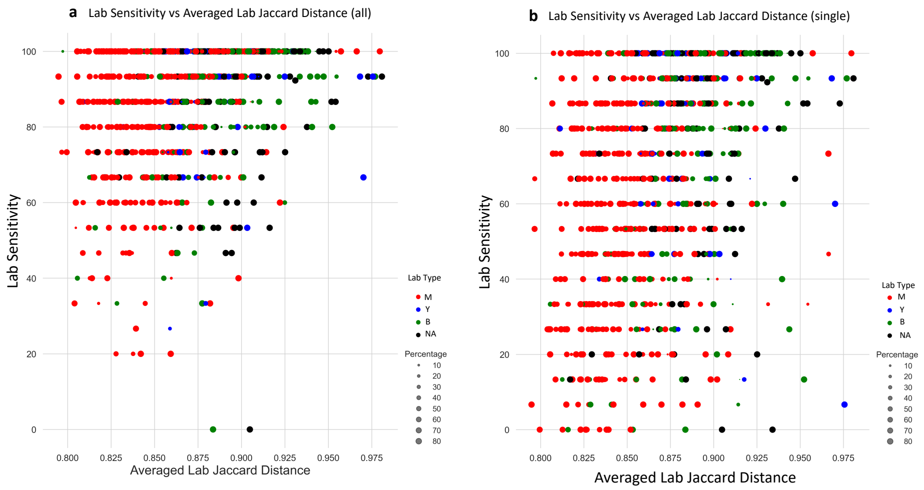


Figure 3: **a** The distribution of lab types displayed over averaged lab Jaccard distances and lab sensitivities (all). Labs with lower lab sensitivities are more likely to have lower averaged lab Jaccard distances as well. It indicates that labs containing less distinct sequences are harder to be correctly identified by $PlasmidHawk$. Since sequences from mammalian labs tend to have high similarities among each others, labs with lab sensitivities below or equal to 60% have higher possibilities to be mammalian labs. **b** The distribution of lab types displayed over averaged lab Jaccard distances and lab sensitivities (single). Among 895 labs, 147 labs have 100% sensitivities. It means that all the plasmids in those labs can be accurately pinpointed to their own source labs.
